# Variational autoencoder for Explainable seizure onset phases detection

**DOI:** 10.1101/2025.09.09.675087

**Authors:** Isaac Capallera, Borja Mercadal, Fabrice Bartolomei, Giulio Ruffini

## Abstract

**Objective:** We present the first deep learning framework for automated, time-resolved, per-channel annotation of ictal and Low-Voltage Fast Activity (LVFA) onsets in stereo electroencephalography (SEEG) recordings of patients with focal epilepsy. To our knowledge, no prior system jointly addresses these tasks on continuous single-channel recordings.

**Approach:** A one-dimensional Variational Autoencoder (VAE) encodes 2-second SEEG segments into a 60-dimensional latent space and classifies them as interictal, ictal, or LVFA via a linear classifier. A postprocessing algorithm converts segment-level probabilities into per-channel onset markers at 0.5-second resolution. The system was trained and evaluated using subject-wise 5-fold cross-validation on 37 patients with manual ictal and LVFA annotations.

**Main results:** At the segment level, the VAE classified the three classes with an average recall of 0.88. At the channel level, it reached an ictal recall of 0.84 (0.91 on Seizure Onset Zone channels) and LVFA recall of 0.74, with median onset latencies of 5.0 s and 0.86 s, respectively. As a seizure detector, the system achieved 99.1 % recall with 1 % false positives. Latent dimensions correlated with physiologically interpretable features (amplitude, band powers, spectral flatness, energy ratio). An ablation study showed that the VAE’s reconstruction objective provides dual benefits over a discriminative encoder-only baseline: improved detection performance and stronger alignment between latent dimensions and these clinically meaningful features.

**Significance:** By providing the first time-resolved per-channel framework for joint ictal and LVFA annotation, this work establishes a robust and explainable platform for automated SEEG analysis with potential to substantially reduce clinician workload during presurgical epilepsy evaluation.

## 1 Introduction

Epilepsy is a neurological disorder characterized by recurrent seizures. Despite multiple available treatments, about 30% of patients live with uncontrolled seizures, which contributes to significant morbidity and a decrease in quality of life.^1, 2^ For patients with drug-resistant epilepsy, intracranial monitoring and surgical resection offer a powerful treatment option. In this context, stereo electroencephalography (SEEG), a technique involving depth electrodes implanted in the brain, is widely used to record electrical activity in the brain and identify seizure onset regions before surgery when direct surgery is not possible without the need of invasive recordings.^3, 4^

Given the invasive nature of SEEG, patients are monitored continuously during multiple days after implantation, leading to massive amounts of data that clinicians review to identify seizure types and onset regions. Analysis and interpretation of EEG/SEEG is significantly labor-intensive and prone to inter-observer variability.^5^ Human observers can easily miss small changes, and agreements on seizure onset presence and timing can vary significantly, even between experts.^6^ This motivates the development of automated seizure detection and annotation systems to assist clinicians. Conventional automated seizure detection methods have relied on signal features on the temporal and frequency domains, and rule-based or classical machine learning algorithms. Such features include Fourier/Wavelet transforms and statistical measures of signal variation.^7–11^ Such approaches, while effective in some settings, often require extensive tuning for each patient, as well as domain expertise for feature selection, and can struggle with generalized performance. They also may produce high false alarm rates or miss non-standard seizure patterns. There is a clear need for more robust, adaptive methods that can learn complex patterns in the data without exhaustive manual feature engineering.

Deep learning has emerged recently as an alternative approach for EEG seizure detection given its ability to learn and generate predictions from raw data. In the last decade, researchers have explored numerous deep neural network architectures, including 1D and 2D Convolutional Neural Networks (CNNs), Recurrent Neural Networks (RNNs), and most recently Transformers, to improve automated epilepsy diagnosis.^12–15^ CNN-based models have achieved widespread success, often by transforming EEG signals into spectrogram images for 2D CNN input, by feeding channel time series into 1D CNNs, or combining both.^16^ These models have demonstrated high accuracy in distinguishing seizures from non-seizure events, sometimes exceeding traditional methods. For example, a CNN applied to short EEG epochs can correctly classify seizure vs. non-seizure with over 98% accuracy on certain datasets.^17^ The performance of deep learning methods, coupled with the lack of need for complex feature engineering, has made them state-of-the-art in several seizure detection benchmarks.

Despite these advances, challenges remain. Standard deep learning classifiers often act as “black boxes” with limited insight into what features of the signal lead to a detection. To tackle this issue, modern methodologies have been explored to provide insights into the decision-making process of neural networks. For instance, techniques such as saliency maps visualize the input regions that most strongly influence the model’s predictions, highlighting the specific EEG patterns or frequency bands that contribute to seizure detection.^18^ There are also studies that use neural networks that can learn class prototypes; ideal examples of seizure activity, comparing EEG segments to these prototypes to make decisions.^19^ These explainability methods not only build trust in the model’s outputs but also can act as a powerful base to guide further refinements in future feature extraction processes for seizure detection.

Autoencoder-based models propose to use unsupervised learning for seizure detection. An autoencoder is a neural network that compresses signals into a fixed, lower-dimensional set of features, known as the latent space, with the assumption that such set of features can fully describe the input data. Then, it uses the latent space to reconstruct the input data with minimum error.^20^ By analyzing this latent space, researchers can identify which signal characteristics are most relevant to the network’s predictions. In a broader sense, latent space analysis allows for the exploration of how different features, such as frequency content, amplitude variations, or temporal patterns, are encoded within the network. This insight provides an important explainability component, offering a window into the decision-making process of the model and facilitating a deeper understanding of the physiological markers associated with seizure activity.

Variational Autoencoders (VAEs) extend this concept by learning a probabilistic latent space. Instead of encoding data to a fixed set of values, it encodes them into a distribution, out of which the latent vectors can be sampled from.^21^ This approach is known to regularize the latent space, improving generalization and leading to a more informative latent space. It has been shown that VAE networks, acting as an anomaly detector with a simple threshold-based decision rule, could detect seizures with near-perfect accuracy on a private EEG dataset.^22^ In that study, 22 of 24 patients achieved 100% seizure detection accuracy. Similarly, Yıldız et al. introduced an unsupervised VAE framework for seizure identification, achieving up to 0.83 AUC in distinguishing seizures from non-seizures on intracranial EEG.^23^ These anomaly-detection frameworks are especially attractive for datasets with very few seizures or highly imbalanced classes. Such methods also make explainability possible via analysis of the latent space.

Other studies have focused on data augmentation and/or semi-supervised learning instead. For instance, You et al. developed a VAE as an anomaly detector that was personalized to each subject. By calibrating the latent space to each patient, they reached a sensitivity of 94.2% at only 0.29 false alarms per hour.^24^ Ganti et al. found that adding synthetically generated SEEG signals using a Generative Adversarial Network to the dataset led to a significant boost in the classification accuracy of their LSTM classifier.^25^ Deep learning models have even been applied to continuous implantable devices: in one study using data from an implanted Neurostimulator (RNS) in 113 patients, a deep seizure classifier achieved around 96% accuracy.^26^ These works illustrate the trend of combining deep learning with clinical EEG/iEEG data, moving toward real-world applicability.

In patients with drug-resistant focal epilepsy, SEEG is a critical component of phase II presurgical evaluation, performed when noninvasive studies are insufficient to precisely localize the epileptogenic zone. SEEG implantation is guided by an a priori hypothesis created from scalp EEG and imaging.^27, 28^ During SEEG review, clinicians manually analyze multiple seizures to determine the exact timing, location, and electrographic morphology of the earliest electrographic changes across implanted channels, as these features are essential for delineating the seizure onset zone (SOZ), distinguishing primary onset from early propagation, and guiding surgical planning.^4, 29^ Among seizure onset patterns, low-voltage fast activity (LVFA) is one of the most common and clinically significant, characterized by low-amplitude, high-frequency activity that often represents one of the earliest electrographic signatures of focal seizure on-set.^29, 30^ LVFA has been widely recognized as a reliable biomarker of the SOZ and has been associated with improved postsurgical seizure reduction, making its identification particularly relevant for epilepsy surgery planning.^29, 31^

Previous VAE-based approaches have successfully utilized anomaly detection for binary seizure identification at the recording level. However, to the best of our knowledge none of them combines three critical features required for presurgical planning: 1) channel-specific annotations, 2) temporal onset markers indicating the precise time when seizure phases begin on each channel, and 3) LVFA detection. In fact, few studies distinguish between standard ictal rhythms and local onset patterns like LVFA.^8, 32, 33^ To address this, we propose a VAE-based deep learning system for automatic SEEG annotation. Trained clinicians conduct SEEG sessions under continuous observation and manually annotate seizures in real time. Thus, our system assumes that seizure-level annotations are available and can be used to extract data segments centered around ictal events. The focus of the proposed method is on automatically labeling the presence and their onset times of ictal and LVFA phases within those segments at the level of single bipolar recordings. The proposed system aims to help clinicians by automatically annotating clinically relevant events in SEEG, thereby reducing the time required for manual SEEG review during the process of identifying seizure onset and propagation zones.

In the present study, we trained a 1D VAE network to classify SEEG segments into ictal, interictal, and LVFA. We then developed a post-processing method to annotate channels based on the VAE predictions. We validated the method using a cross-validation analysis on 37 patients. We also analyzed the explainability of the network’s predictions by comparing the latent dimensions with SEEG features. The results show competitive performance with state-of-the-art and highlight how specific latent dimensions of the VAE correspond to interpretable SEEG electrical phenomena.

While other, more powerful generative architectures exist, such as GANs or diffusion models, they are more focused on generation rather than representation. That is, even though the outputs have a strong resemblance to real data, their internal processes are much harder to interpret and correlate with clinical SEEG features than the VAE’s latent space. Since the main scope of this study was to exploit the inner latent spaces of a network to distinguish between seizure phases, with reconstruction fidelity being an auxiliary task, the choice of the VAE as architecure became the most practical and effective choice. Rather than just learning how to recreate brain waves, the VAE acts like a smart compressor. By forcing the complex SEEG signals through a narrow bottleneck, it organizes the underlying data into a structured space. This allows us to clearly observe how different seizure phases naturally group together, providing an explainable window into clinical practice that purely generative models tend to obscure. In the following sections, we describe our approach for using a VAE based framework to automatically annotate ictal and LVFA periods on SEEG. We detail the architecture and training strategy of the VAE, the data processing steps, a postprocessing algorithm to aggregate the classifications of SEEG segments to a channel level prediction, the system’s subject-wise cross validation strategy. Finally, we describe how we will assess the explainability of the full system.

## 2 Methods

The system is designed with the main objective of processing an SEEG recording and returning markers indicating the start of ictal and/or LVFA events, when present, in the different channels. A second objective is the detection of seizures in any given recording based on its individual channels. Throughout this study, the term *channel* will be used to refer to the differential SEEG recordings between adjacent contacts (bipolar montage). The system is meant to act as an annotating tool as depicted in 1A and is designed in multiple parts: first, the SEEG recordings in our dataset are annotated and split into 2-second segments to train a VAE network coupled with a linear classifier, converting the VAE into a supervised learning network, where the VAE learns a latent space that enables both reconstruction of the segments with minimal error and high classification performance, as depicted in Figure 1B. Then, a postprocessing threshold is adjusted for an algorithm that uses the AUC of the probabilities of all segments in a channel for classification, allowing the placement of markers on a fine-grained temporal scale. Our 37-subject dataset is divided into five different folds for cross-validation, allowing us to train the VAE, adjust the postprocessing algorithm, and test the full system with all available data, ensuring subject independence on the three processes. The data split strategy is illustrated in Figure 1D. Finally, the system is used as a seizure detector per channel and per recording.

**Figure 1:**
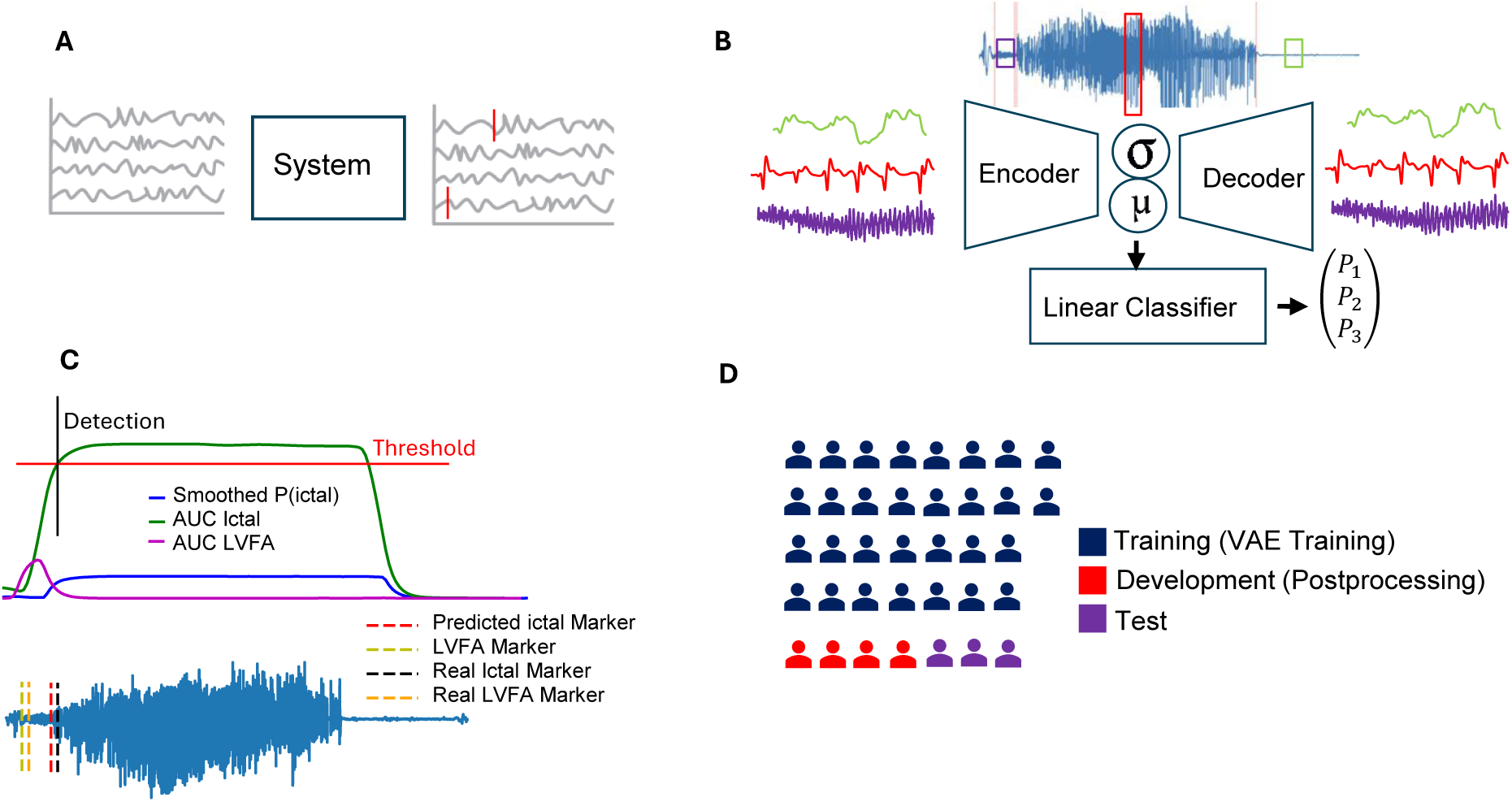
Overview of the proposed methodology. **A**: Functionality of the full system, where individual channels of SEEG recordings can be annotated automatically. **B**: SEEG channels are annotated and split into segments of interictal, ictal and low voltage fast activity (LVFA) classes, which are used to train the variational autoencoder (VAE) and obtain reconstructions and class probabilities per segment. **C**: A postprocessing algorithm is used to convert segment probabilities of SEEG channels into markers. The area under the curve (AUC) of the probability segments is computed for the ictal and LVFA classes, placing a marker where the AUC surpasses a tunable threshold. **D**: The 37 available subjects are split into five folds for cross-validation, with each fold containing separate training, development, and test sets.

The results are assessed at multiple levels. Firstly, we analyze the classification performance of the VAE, trained on the 2-second fragments. Secondly, the full channel annotation performance after the postprocessing. Finally, as a secondary objective, we evaluate the capability of the full system to recognize seizure recordings based on the prediction of all channels per recording.

### 2.1 Data extraction

In the present study, data from 37 different subjects was used from two different datasets. In terms of available seizures, our dataset is comprised of 108 seizures, with each subject containing 3 available seizures on average. A summary of the subject demographics and clinical characteristics can be found in Table 1 and a detailed description of the dataset can be found in the Supplementary table S1. The cohort includes 11 patients with mesial temporal lobe epilepsy, 13 with temporal lobe epilepsy and 13 with extratemporal epilepsy (frontal, parietal, occipital). LVFA patterns were present in 18/37 subjects (49%), with heterogeneous manifestation: some subjects showed LVFA in multiple channels across seizures, while others showed LVFA in only a subset of SOZ channels. The number of available channels per subject ranges from 36 to 218, mean 119*±*45. SOZ size ranged from 3 to 25 channels per subject (mean 10*±*6). This heterogeneity reflects real-world clinical diversity and tests the model’s ability to generalize across different seizure onset patterns, electrode configurations, and epilepsy types

**Table 1:** Dataset summary with subject demographics and clinical characteristics. Values shown as mean*±*standard deviation for continuous variables or count (percentage) for categorical variables.

| Characteristic | GALVANI (n=13) | HUP (n=24) | Total (n=37) |
| --- | --- | --- | --- |
| **Age** | 33 ± 11 | 33 ± 10 | 33 ± 10 |
| **Gender** | 5F/8M | 9F/15M | 14F/23M |
| **Epilepsy type** |  |  |  |
| Temporal | 7 | 6 | 13 (35%) |
| Mesial Temporal | 0 | 11 | 11 (30%) |
| Frontal | 2 | 4 | 6 (16%) |
| Other | 4 | 3 | 7 (19%) |
| **MRI Lesion** | 3 | 12 | 15 (41%) |
| **Channels per subject** | 153 ± 34 | 100 ± 38 | 119 ± 45 |
| **Seizures per subject** | 2 ± 1 | 3 ± 1 | 3 ± 1 |
| **SOZ size (Channels)** | 11 ± 5 | 9 ± 6 | 10 ± 6 |
| **LVFA presence** | 9 | 9 | 18 (49%) |

Most of the data was obtained from the publicly available iEEG HUP dataset.^34^ This dataset includes recordings from 58 subjects who underwent ECOG or SEEG monitoring as part of surgical evaluation for drug-resistant epilepsy at the Hospital of the University of Pennsylvania. For this study, only subjects with SEEG data available in a bipolar montage were selected, which reduced the number of available subjects to 24. The dataset contains recordings during the ictal phase of seizures, as well as interictal recordings. The former was used as data to train the system, while the latter was used as validation data to assess false positive detections. The dataset includes clinician annotations indicating the channels belonging to the seizure onset zone, as well as time stamps indicating the global start and end of each of the available seizures.

The second dataset consists of 13 subjects with refractory focal epilepsy from the GALVANI project, extracted from different clinical trials (NCT04782869, NCT05250713).^35–37^ A list of channels belonging to the epileptogenic and propagation zones, as well as annotations indicating the seizure onset times, was provided by clinicians for each seizure.

### 2.2 Data Processing

Segments containing the seizure events were extracted from all the 108 total recordings. segments with a duration approximately twice that of the ictal period were extracted to maintain a balance between ictal and interictal activity. A high-pass filter at 0.5 Hz and a notch filter (50 Hz for Galvani data and 60 Hz for the HUP data) were applied to remove powerline interference. Then, recordings were resampled to a common sampling rate of 512 Hz. Channels with big movement artifacts were rejected, and each channel was standardized using z-score normalization.

Seizure onset and offset annotations were provided by the clinicians in channels in the EZ and did not include further detailed labeling at the individual channel level. We manually refined these annotations to produce channel-specific onset markers in order to enable channel-specific analysis. For consistency, the start of an ictal phase was marked by following these guidelines:^38^

1. Presence of a sudden rise in amplitude in the SEEG signal.

2. Such activity is rhythmic and happens mostly in delta, theta, alpha, or beta bands in the Power Spectral Density (PSD) signal.

In contrast, no clinical annotations for LVFA were available in the datasets. Therefore, the presence of LVFA events was manually asserted on channels strictly labeled by the clinicians as part of the EZ or propagation zone. Given the subjective nature of the task, to mitigate uncertainty and inter-observer variability a consensus based labeling process was performed: annotations were primarly performed by one author and reviewed by a second author. Furthermore, the existing clinician annotated ictal marker served as an anchor, restricting search for the LVFA to a time window shortly preceding the ictal start. Finally, to ensure consistency, the following rules were applied to the annotation of LVFA:^29^

1. Electrical activity must be lower or at a similar amplitude to baseline SEEG activity.
2. There is activity on the PSD in the high beta band or higher, either in a broadband or in the form of a chirp.

The time series of the different channels were segmented into 2-second segments. Data augmentation was performed on each training set using a sliding window method to minimize class imbalance, especially around the LVFA class. Namely, while non-overlapping segments were used for the interictal class, sliding windows of 1 and 0.1 seconds were used to extract segments belonging to the ictal and LVFA classes, respectively. In addition, amplitude scaling was used on all segments at scales of 0.5 and 1.5.

For the extraction of interictal recordings, a maximum of 3 minutes of recording were extracted for each available recording. If less than 3 minutes of interictal data were available in the recording, all interictal data was extracted instead, ensuring no overlap with the segments extracted for ictal data. While 108 total recordings were available, only 102 interictal recordings were extracted, since 6 recordings contained interictal periods deemed too short.

### 2.3 VAE architecture and training

A symmetrical architecture for the encoder and decoder was adopted for the VAE, with construction performed using the TensorFlow library in Python. They both consist of five 1D Convolutional layers with progressive temporal downsampling followed by two Bi-LSTM layers, which are known to extract relevant features from time series data. The mean and standard deviation that parametrize the posterior distribution of the VAE were modeled using linear layers, with the latent space size being 60 dimensions. A single linear layer is used for the classifier, with the means of the generated distributions taken as its input. A normal Gaussian was selected as the prior distribution, as in most VAE applications. All per-layer hyperparameters are further detailed in Table 2.

**Table 2:** Summary of all the layers in the VAE architecture and its hyperparameters

| Layer Name | Channels/Units | Kernel Size | Padding | Activation |
| --- | --- | --- | --- | --- |
| **Encoder – Convolutional Layers** |  |  |  |  |
| 1D Convolution 1 | 32 | 10 | same | ReLU |
| Max Pooling 1 | – | 2 | – | – |
| 1D Convolution 2 | 64 | 10 | same | ReLU |
| Max Pooling 2 | – | 2 | – | – |
| 1D Convolution 3 | 128 | 10 | same | ReLU |
| Max Pooling 3 | – | 2 | – | – |
| 1D Convolution 4 | 256 | 10 | same | ReLU |
| 1D Convolution 5 | 512 | 10 | same | ReLU |
| **Encoder – LSTM Layers** |  |  |  |  |
| BiLSTM1 | 128 | – | – | Tanh |
| BiLSTM2 | 256 | – | – | Tanh |
| **Latent Space** |  |  |  |  |
| Dense $z_\mu$ | 60 | – | – | Linear |
| Dense $z_{\log \sigma^2}$ | 60 | – | – | Linear |
| Dense (Linear Classifier) | 3 | – | – | Linear |
| **Decoder – LSTM Layers** |  |  |  |  |
| Dense | 65536 | – | – | ReLU |
| Reshape to $(128 \times 512)$ | – | – | – | – |
| BiLSTM1 | 256 | – | – | Tanh |
| BiLSTM2 | 128 | – | – | Tanh |
| **Decoder – Transposed Convolutional Layers** |  |  |  |  |
| 1D Transposed Convolution 1 | 512 | 10 | same | ReLU |
| Up Sampling 1 | – | 2 | – | – |
| 1D Transposed Convolution 2 | 256 | 10 | same | ReLU |
| Up Sampling 2 | – | 2 | – | – |
| 1D Transposed Convolution 3 | 128 | 10 | same | ReLU |
| Up Sampling 3 | – | 2 | – | – |
| 1D Transposed Convolution 4 | 64 | 10 | same | ReLU |
| 1D Transposed Convolution 5 | 32 | 10 | same | ReLU |
| Output Transposed Convolution | 1 | 10 | same | Linear |

This architecture was designed following established principles for deep learning on physiological time series. We adopted a CNN-LSTM combination commonly used for EEG/SEEG analysis.^39–42^ The five convolutional layers with progressive channel expansion (32*→*512) provide multi-scale feature extraction while the two BiLSTM layers (128, 256 units) model the long range temporal dependencies. An exhaustive architecture search was not performed due to computational constraints. However, our design is well-motivated by prior work demonstrating that CNN-LSTM combinations effectively capture both local patterns (via convolution) and temporal dynamics (via recurrence) in physiological signals.^39–42^ The specific layer counts and channel sizes represent a reasonable balance between model capacity and the risk of overfitting given our dataset size.

The latent dimension size of 60 was selected through systematic evaluation of the reconstruction ability for various (Supplementary Figure S1). Reconstruction loss decreases with increasing latent dimensions, with diminishing returns beyond 60 (Supplementary Figure S1). Visual inspection confirmed that 60 dimensions adequately reconstruct the dominant patterns in ictal activity, while LVFA reconstruction remains challenging across all dimensions. The latent size choice was also made to balance reconstruction quality with model compactness, favoring generalization and computational efficiency. Larger dimensions offered marginal reconstruction gains at the cost of increased complexity.

The VAE was trained in a supervised setting, with access to labels leveraged to encourage encoding of class-relevant features in the latent space. Three components, reconstruction error, KL divergence, and categorical cross-entropy for classification, are combined in the VAE’s loss function, which is detailed as follows:

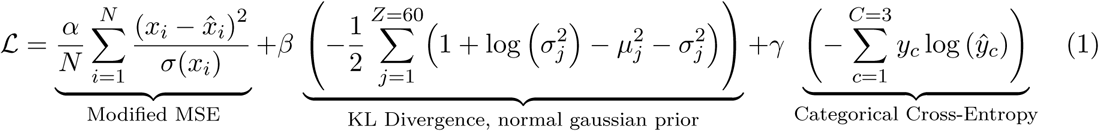

The MSE equation was slightly modified to penalize reconstruction of high-amplitude signals in favor of low-amplitude ones by dividing it by the segment’s standard deviation. The weight of each component of the loss function was adjusted heuristically at *α* = 2000, *β* = 0.01, and *γ* = 110. The network was trained for 50 epochs using the AdamW optimizer with a base learning rate of 0.001. A learning rate scheduler was used to reduce the base learning rate when the validation loss did not improve for ten epochs, which helps avoid local minima. All training activity was performed in a Nvidia RTX A-6000 GPU.

### 2.4 Postprocessing

For whole-channel seizure annotation, Each SEEG channel is segmented into overlapping 2-second segments using a 0.5-second sliding window. Class probabilities for every segment are obtained using the encoder and classifier of the VAE network. An exponential smoothing is applied to the ictal/LVFA probability sequences to reduce sudden fluctuations and stabilize the temporal profile of the probabilities. The AUC of the smoothed probabilities is computed using a moving window of 8 and 3 segments for the ictal and LVFA classes, respectively. To select the window sizes, we looked at the shortest annotated ictal and LVFA periods available in the dataset, which were of 4 and 1.5 seconds, respectively. Since the sliding window of the postprocessing algorithm is 0.5, we set the window size to be just big enough to cover the shortest seizure.

Ictal and LVFA events are detected using a threshold-based approach applied to the AUC. The onset time is assigned to the first segment where the AUC surpasses the threshold. For LVFA detection only, an additional constraint is imposed: the LVFA marker is only considered valid if an ictal marker is also present in the channel, and if the predicted LVFA occurs prior to the ictal marker.

Following this pipeline, a single marker per class is obtained per channel, indicating the time in the recording where the Ictal/LVFA event begins

### 2.5 5-Fold Cross Validation and evaluation

A subject-wise 5-fold cross validation was implemented to ensure the robustness and generalization of the proposed VAE-based framework. The complete dataset, comprising SEEG recordings from 37 subjects, was partitioned into 5 mutually exclusive folds so that each subject appears in exactly one fold. In each cross-validation iteration, one fold is held out as the validation set, while the remaining four folds are used as the training set. It is guaranteed by this procedure that training and evaluation are performed on independent recordings (i.e. from different subjects). Table 3 details the distribution of segments per class across the five folds, highlighting the strong class imbalance in the dataset. even after the augmentation methods described in section 2.2:

**Table 3:**
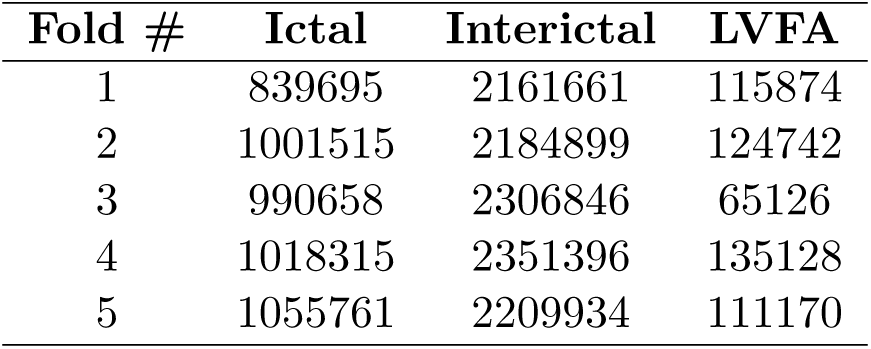
Segment counts for each class per fold after data augmentation. The table highlights the class imbalance: the Interictal class is the most prevalent (approximately. 67%), while the LVFA class is in a strong minority (*<*4% of the total dataset, approximately).

| **Fold #** | **Ictal** | **Interictal** | **LVFA** |
| --- | --- | --- | --- |
| 1 | 839695 | 2161661 | 115874 |
| 2 | 1001515 | 2184899 | 124742 |
| 3 | 990658 | 2306846 | 65126 |
| 4 | 1018315 | 2351396 | 135128 |
| 5 | 1055761 | 2209934 | 111170 |

For each iteration, the training set, after data processing, is used to train the VAE. The validation fold is further subdivided into two independent subsets: a development set and a test set. The development set is used to tune the detection threshold employed in the postprocessing algorithm.

The optimal threshold was determined through a parameter sweep. For the ictal class, this tuning was performed within each fold: the optimal threshold was selected on the development subset based on achieving over 90% precision while maximizing detection accuracy and then evaluated on the test subset.

For the LVFA class, however, the number of subjects exhibiting this pattern was insufficiently distributed across the folds. Therefore, a global pooling approach was adopted: all LVFA-positive subjects from the entire dataset were aggregated into a single dataset for threshold tuning. A more stable and representative threshold for LVFA detection was obtained as a result.

Finally, the subjects in the development and test subsets are interchanged. Namely, after tuning the detection threshold on one subset and evaluating on the other, the process is then repeated in reverse. Every subject in the dataset is served as an independent test case across the cross-validation strategy by this two-pass approach. Thus, potential bias from data partitioning is reduced, and a more robust and exhaustive assessment of the system’s performance is achieved.

The system is evaluated at multiple scales. Firstly, we evaluate the classification and reconstruction performance of the VAE alone, which operates at the 2-second segment level. Performance metrics, such as recall, precision, F1 score and MSE for reconstruction, are computed at this stage.

The primary object of the proposed system, however, is to serve as an automatic annotation tool for SEEG recordings at the channel level. As such, channel-level detection is the primary focus of the performance evaluation. Namely, the automatic annotations on each channel are compared with the manual labels. Based on that, performance metrics, including accuracy, precision, recall, specificity, and F1 score, are computed for each subject. Additionally, the temporal accuracy is quantified by comparing the detection times on each channel with the manually annotated onsets. False positive detection using the interictal channel recordings of all subjects is also assessed, with specificity across all channels per subject evaluated in the same fashion as channel-level detection.

Finally, the performance of the system as a seizure detection tool is also assessed. In this case, a seizure is deemed as detected if at least one channel registers the event. Seizure detection is evaluated as the percentage of correctly detected seizures. Similarly, recordings with no ictal activity are used to determine the false positive rate as a seizure detector.

## 3 Results

### 3.1 System Performance Assessment

We trained the VAE network in each cross-validation fold, resulting in five separately optimized models. Before integrating the VAE into the full seizure annotation pipeline, we evaluated its performance to confirm consistency across folds and to verify that the latent space effectively captures features relevant to seizure classification. Figure 2 presents a summary of performance at reconstructing and classifying the 2-second segments evaluated on the validation set, thereby demonstrating both the consistency and reliability of the VAE training. This per-fold evaluation confirms that our model not only learns to accurately reconstruct the input signals but also successfully captures class-dependent features in its latent space that are critical for discriminating between ictal, interictal, and LVFA segments.

**Figure 2:**
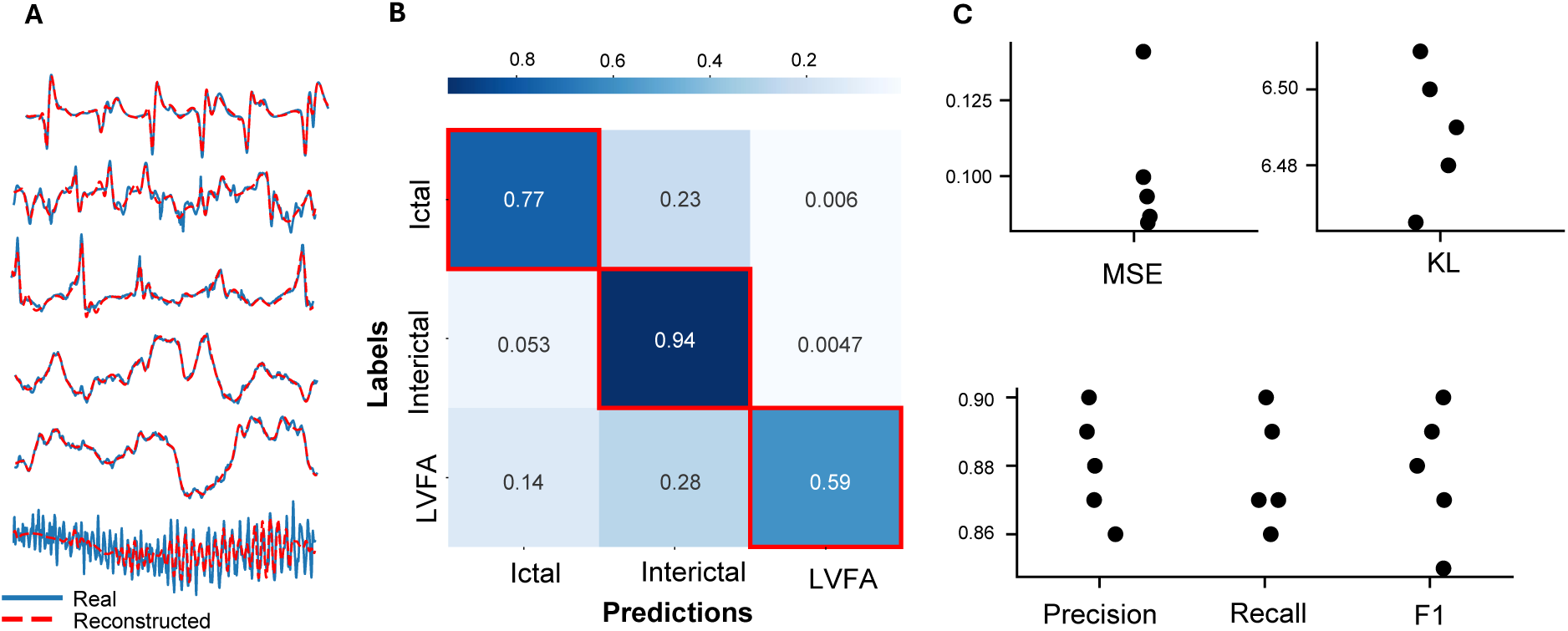
**A:** Reconstruction examples from different segments of the three classes **B:** Confusion matrix indicating the variational autoencoder classification performance, normalized per class **C:** Metric numbers for mean squared error (MES), Kullback-Leibler (KL) divergence and Classification Precision, Recall and F1 scores for the five different folds. The classification metrics were obtained per fold as a weighted average of the three classes.

Figure 2A displays examples of signals reconstructed by the VAE. We can observe that while low frequency components are accurately reconstructed, high frequency information, mainly observed in LVFA segments, is primarily under-reconstructed. This discrepancy reflects a phenomenon known as spectral bias in neural networks,^43^ where models based on gradient optimization, such as neural networks, pushes the network to first fit the dominant, slowly varying structure of the signal, leaving the finer, high-frequency details as residuals that are learned only slowly or not at all. In the context of autoencoders, spectral bias manifests as a higher reconstruction error where sharp, high frequency content dominates.

Figure 2B provides the average confusion matrix over the five cross-validation folds at the segment level. The model is able to correctly classify 77% of ictal segments and 94% of interictal segments, however, only 59% of LVFA segments are correctly identified. The most common errors arise from ictal and LVFA segments being confused with interictal (23% of ictal segments and 28% of LVFA segments), particularly reflecting the difficulty of distinguishing high-frequency bursts from background activity. A smaller fraction of LVFA segments (14%) are mislabeled as ictal, and a modest 5.3% of interictal segments are confused for ictal.

Finally, we present the reconstruction error, the KL divergence and the classification performance of the VAE in Figure 2C. The results further highlight that the VAE achieves high classification performance at the segment level, with precision, recall, and F1 scores reaching approximately 0.88 on average. In parallel, the reconstruction error and KL divergence values indicate a balanced trade-off between reconstruction fidelity and latent space regularization, confirming that the network is learning a meaningful, structured representation of the data. These consistent results across folds underscore the robustness of our approach and provide a strong foundation for the subsequent channel-level and seizure-level analyses.

After analyzing the performance on the VAE alone, we coupled it with our Postprocessing algorithm to evaluate the performance of the whole system, that is, the performance as a whole channel annotator. We adjusted the detection thresholds following the methodology in section 2.4. We summarize the optimal thresholds in table 4. Notably, the optimal threshold values across sets lie within a narrow range, indicating the robustness of the system to the subset of subjects used for tuning. This consistency also highlights the potential of the system for generalization to unseen data. For the LVFA class, due to the limited number of examples, a single global threshold was computed, yielding a value of 1.08.

**Table 4:**
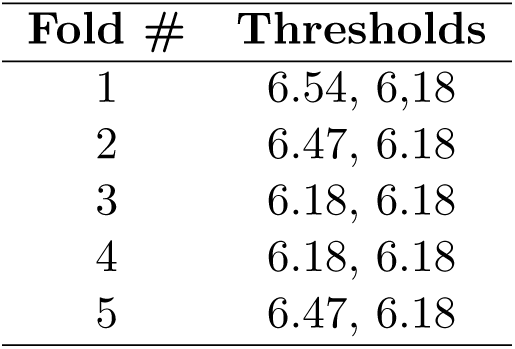
Optimal threshold values obtained in each fold. Note that two values are reported per fold due to the two-pass approach used within the 5-fold cross-validation procedure (see section 2.5)

With the threshold properly tuned, we used the postprocessing algorithm to annotate the recordings of all subjects. The dataset contained 108 seizures in total.

We analyzed the performance of our system at the individual channel level. As such, presence of manual and predicted markers are tracked on each channel per subject. We evaluate the detection in a subject-wise manner, computing the reported metrics using all its channels and ictal recordings, comparing the presence of automatic annotations against human labels like a traditional, cascaded, binary classification problem. In the case of time difference, we used the values of the manual and predicted markers to obtain the difference in seconds between the predicted and manual markers. We present the results of ictal detection performance assessment in Figure 3A. The subject-wise evaluation revealed an average and standard deviation accuracy of 0.92*±*0.09, with a median of 0.95; precision of 0.9*±*0.1, with a median of 0.95; recall of 0.8*±*0.2, with a median of 0.95; specificity of 0.9*±*0.1, with a median of 0.95; and an F1 score of 0.85*±*0.17, with a median of 0.92. Additionally, when focusing on channels that were specifically labeled as part of the seizure onset zone, the average recall increased to 0.9*±*0.2, with a median of 1.

**Figure 3:**
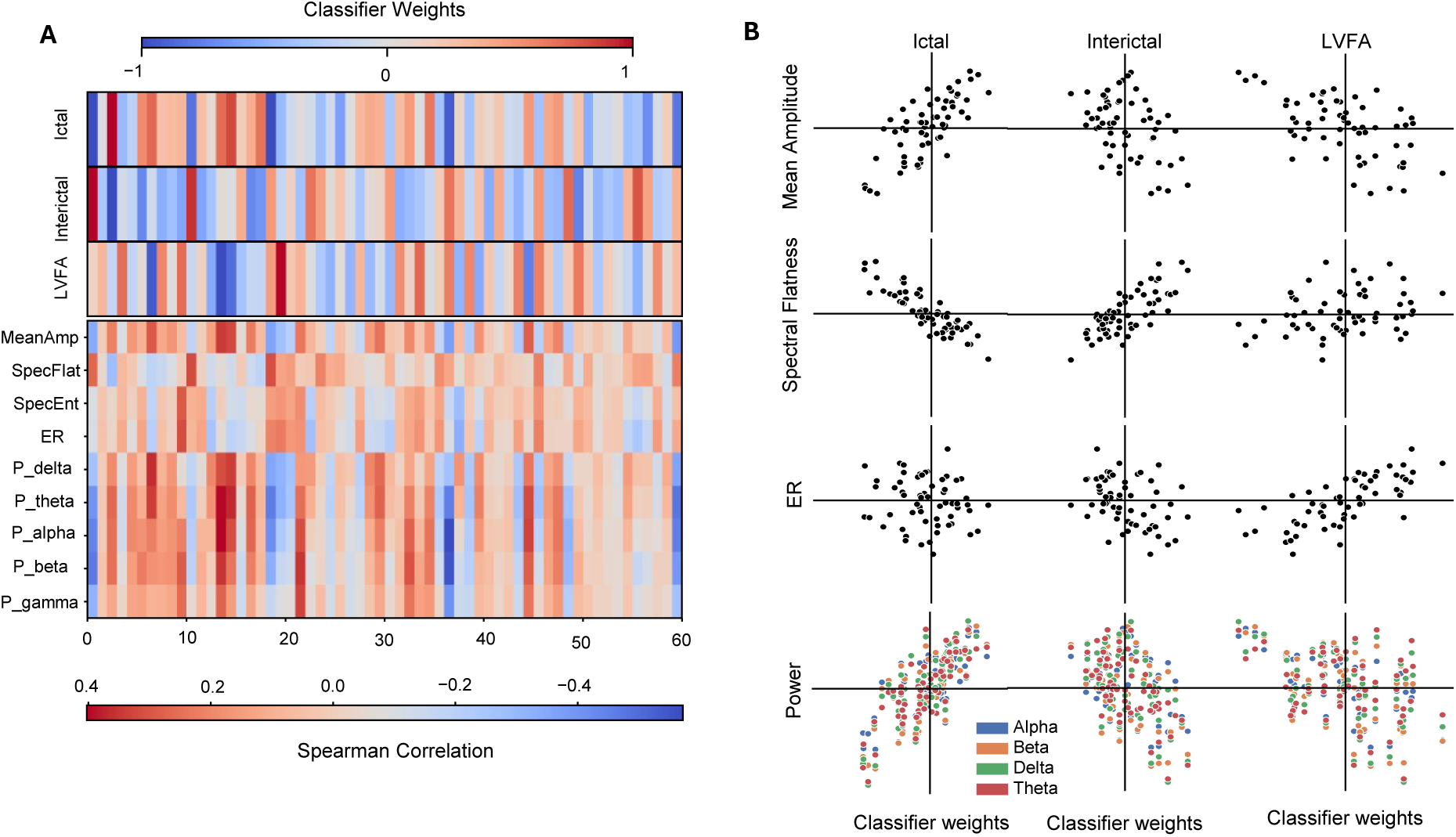
**A:** Metrics for the ictal detection, evaluated as an average of all channels per subject. Individual points indicate performance on each subject. Note that recall is evaluated across all channels as well as on the channels labeled as part of the seizure onset zone (SOZ) by clinicians. **B:** Metrics for the low voltage fast activity (LVFA) detection. **C:** Histogram of the time lag, in seconds, between the real and predicted ictal annotations of the whole dataset, clipped to 50 seconds before and after the real marker. **D:** Histogram of the time lag, in seconds, between the real and predicted LVFA annotations of the whole dataset, clipped to 15 seconds before and after the real marker.

As evidenced by Figure 3A, detection performance is high overall on individual subjects, especially at seizure onset channels, where ictal activity is more evident. However, a few subjects display a poor detection performance. Upon inspection of the recordings of these subjects, most detection failures can be attributed to one or more of the following signal characteristics: 1) Short ictal duration (10 seconds or less) compared to other subjects; 2) low-amplitude ictal activity, leading to misclassification as interictal; and 3) dominant high frequency components, leading to misclassification as LVFA.

In terms of LVFA detection, the evaluation methodology was slightly different than the ictal period, since, unlike the ictal class, we adjusted a single global threshold for all subjects. As depicted in Figure 3B, we observe an average and standard deviation accuracy across all subjects of 0.98*±*0.08, with a median of 0.99; precision of 0.8*±*0.2, with a median of 0.91; recall of 0.7*±*0.3, with a median 0.87; specificity of 0.97*±*0.08, with a median of 0.99; and F1 score of 0.7*±*0.3, with a median of 0.80. Such performance is due to a combination of causes, the main ones being:

- The aforementioned spectral bias affecting our network, causing high frequency components in LVFA to be harder to learn.
- Manual annotations for LVFA were not explicitly provided by clinicians, but instead determined by us following concrete rules. Additionally, there are less LVFA events compared to the other classes, causing class imbalance that our system struggles to deal with.
- Ictal activity was not detected on the channel preventing the system from annotating the LVFA onset
- False positive detections due to unrelated high-frequency activity.

Besides the detection performance, we also evaluated the temporal accuracy of the system’s annotations by calculating the difference in time compared to manual annotations. We’ve condensed all the differences from all channels and subjects in a histogram in Figure 3C, showing that most of the differences are centered around zero. The average time difference for the ictal class was 9.80 seconds, with a median of 5.02 seconds. We observe a similar pattern when analyzing the time differences in the LVFA annotations in 3D; however, given the natural shorter duration of the LVFA period with respect to the ictal duration, the range of differences is shorter. The average time difference in the whole dataset was 2.04 seconds, with a median of 0.86 seconds.

The results show that while many ictal onsets are detected nearly synchronously with human labels, there is still variability in detection latency. Specifically, in the ictal phase, most of the latency is originated by detecting the ictal onset late, rather than wrongfully detecting interictal spikes as part of the ictal phase.

In contrast, in the LVFA class, detection time offsets are smaller and equally distributed in both signs (i.e. there is a similar amount of early and late detections). However, there were a few outliers 30 to 60 seconds earlier than the real annotations, suggesting that high-frequency artifacts during the interictal period were incorrectly detected as LVFA.

Additionally, even though this was not the main scope of the study, we evaluated the performance of the system for seizure detection.

To ensure consistency, we considered a seizure as successfully detected if it was correctly marked on at least one channel. Figure 4 depicts the performance as a seizure detector in the form of a binary problem between seizure and no seizure presence evaluated on all available recordings, using all their channels. Out of the 108 ictal recordings, 107 seizures were identified, yielding a seizure detection recall of 99.1% and a false positive rate of 1%. In terms of false positives 5 out of 102 interictal recordings had at least one channel with an incorrect ictal marker prediction, with an average of 2 incorrect channels per recording, yielding a false negative rate of 4.5%. The low amount of false positives in interictal recordings reinforces the robustness of the system, proving that the system only struggles to classify ambiguous cases.

**Figure 4:**
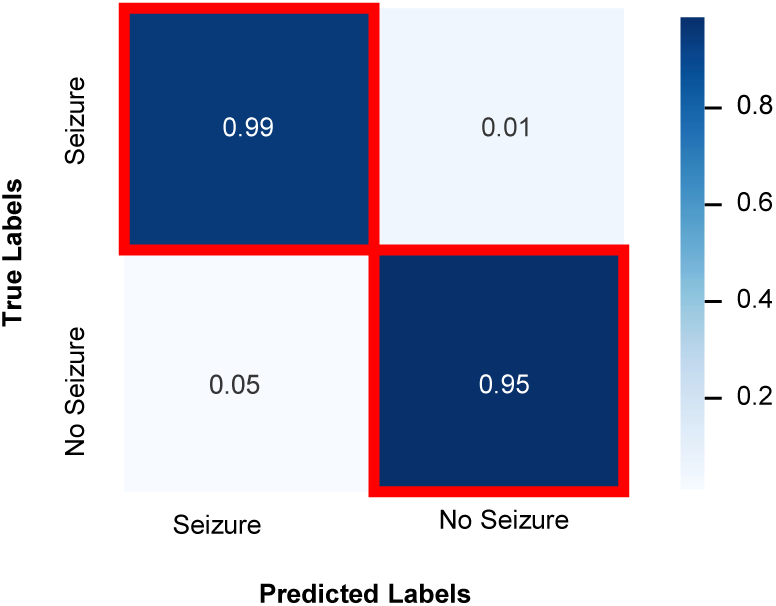
Normalized confusion matrix of the classification performance on entire seizure and non-seizure recordings. The seizure detection task is a binary classification problem over the entire SEEG recording and all its channels. Our dataset had a total of 108 ictal and 102 interictal recordings

### 3.2 Latent space analysis

A key motivation for using VAEs in clinical deep learning applications is their potential for increased explainability compared to other network architectures. Unlike conventional black-box models, VAEs learn a structured latent space representation of the data that can be analyzed. By doing so, it is possible to investigate which aspects of the input data are more relevant for the classification task. To evaluate this explainability in our system, we performed a detailed analysis of the learned latent representations after retraining the VAE on the entire dataset.

The three classes in our classification task are typically characterized as follows:

1. Ictal activity is characterized by high amplitude and a narrow band activity in the delta, theta or alpha bands.
2. LVFA activity presents the opposite behavior, characterized by low amplitude activity and spectral power dominated by a higher frequency range, which can be depicted by a higher energy ratio.
3. Interictal activity is generally less rhythmic (i.e. broadband) associated with a flatter spectral profile compared to ictal and LVFA activity. In terms of amplitude, interictal activity is typically characterized by a lower amplitude than ictal activity.

To investigate which signal features are encoded in the latent space, we analyzed the correlation between the latent space dimensions and a set of spectral features that can putatively be used to discriminate between the different onset phases. The selected features were:

1. Mean amplitude, defined as the average of the absolute values of the signal over a time window.
2. Frequency power at different frequency bands. The power in specific frequency bands is calculated by filtering the power spectral density signal to the band of interest, computing the squared magnitude, and then averaging over the window. The selected bands where: Delta (0.5–4 Hz), theta (4–8 Hz), alpha (8–12 Hz), beta (12–30 Hz), and gamma bands (30–120 Hz).
3. Spectral entropy: It measures the complexity/randomness of the spectrum, computed as . Signals with broadband activity, such as interictal segments will have high entropy while ictal and LVFA segments, which are focused at more specific frequencies, will have low entropy.
4. Spectral flatness: Similarly, it measures the presence of a dominant frequency, computed as . Ictal and LVFA segments will have low flatness, while interictal segments will have high segments.
5. Energy ratio, defined as the power at beta and gamma bands divided by the power at theta and alpha bands. This metric has been previously used to detect onsets at high frequency.^11^

For each 2-second SEEG segment, we obtained its associated vector in the latent space by passing it through the encoder and evaluating the features described. We then computed the Spearman correlation coefficient between the values of the signal features and the latent space dimensions across the entire dataset. In addition, we examined the weights of the linear classifier applied to the latent space, which quantify the contribution of each latent dimension to the prediction of each class. By jointly analyzing the classifier weights and the feature correlations, we can identify which signal features are more relevant to the classification task according to the network.

Figure 5 summarizes the results of this correlational analysis. Figure 5A shows the weights of the linear classifier alongside the Spearman correlation values between the latent dimensions and the selected signal features. Interestingly, the strongest positive and negative weights of the different classes are seen in dimensions that correlate/anti-correlate with the features expected for that class. To further analyze this correlation, we present the same results in scatter plots (Figure 5B).

**Figure 5:**
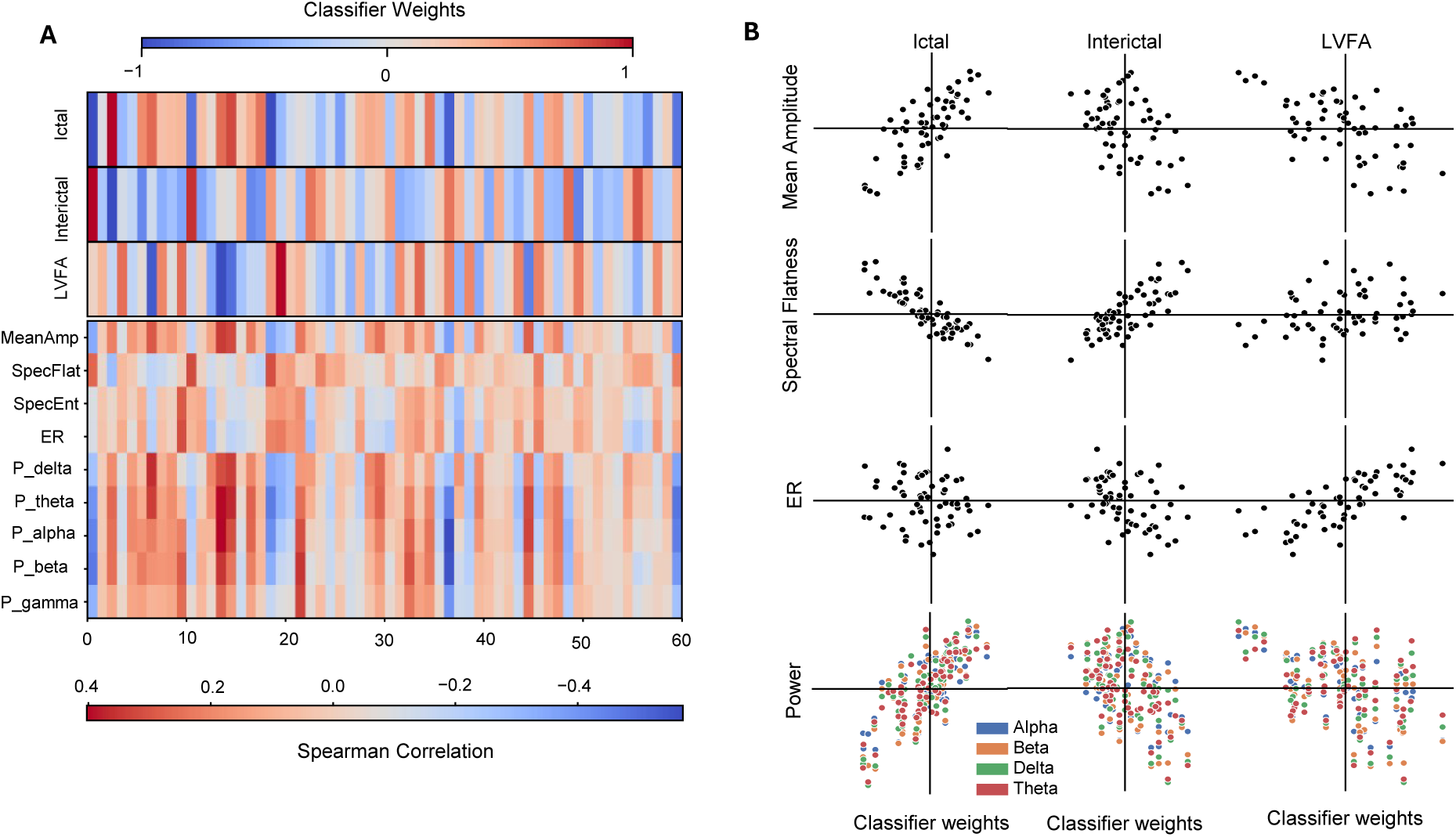
Explainability analysis of the variational autoencoder-based classification. **A**: Heatmap of normalized classifier weights for each latent dimension and class (ictal, interictal, LVFA), along with the Spearman correlations between each latent dimension and several SEEG features. The color scales indicate the direction and magnitude of the weights or correlations. **B**: Scatter plots compare each dimension’s classifier weight (x-axis) to its correlation with selected features (y-axis).

For the ictal class, latent dimensions that are strongly correlated with mean amplitude and with power in the delta, theta, and alpha bands also exhibit the highest classifier weights. In addition, while no correlation is seen between the weights and the ER, the weights are negatively correlated with the spectral flatness. This finding aligns with clinical expectations, as ictal events are typically characterized by an increase in amplitude and narrow-band rhythmic activity in these frequency ranges. Furthermore, dimensions that display anti-correlations with these same features contribute negatively to ictal predictions, indicating that they play a role in mitigating false-positive detections.

In contrast, for the LVFA class, the weights display a negative correlation with the amplitude and the power in the low frequency bands. In addition, the results reveal that latent dimensions with high positive correlations to the energy ratio are particularly relevant for the classifier. This suggests that, despite the spectral bias, the model is effectively recognizing the signature high-frequency, low-amplitude patterns that define LVFA.

Finally, for the interictal class, the most relevant latent dimensions show a strong correlation with spectral flatness, reflecting the absence of dominant frequency components, which is characteristic of normal or background brain activity. Furthermore, the power in the different bands and the amplitude are negatively correlated with the classifier weights. These findings are aligned with the clinical description of interictal activity.

This analysis not only highlights the model’s ability to encode physiologically meaningful features but also provides valuable insights into how these features contribute to the classification of different seizure onset patterns. This level of explainability is crucial for building trust in deep learning applications, an essential requirement for their deployment in clinical settings.

### 3.3 Ablation study: Contribution of Reconstruction Objective

To evaluate whether the VAE’s reconstruction objective contributes to detection performance and latent space explainability beyond the discriminative encoder alone, we conducted an ablation study comparing our full VAE model against an encoder-only architecture.

We trained a deterministic encoder-only model consisting of the identical encoder architecture followed by a linear classifier operating directly on the latent vector representation, without the probabilistic sampling, decoder, or reconstruction loss components of the VAE. This model was trained exclusively on the categorical cross-entropy classification loss. We applied the same 5-fold cross-validation scheme, including identical postprocessing algorithms and threshold tuning procedures as those used with the VAE.

Figure 6A presents the channel-level detection performance metrics for both models across all subjects.

**Figure 6:**
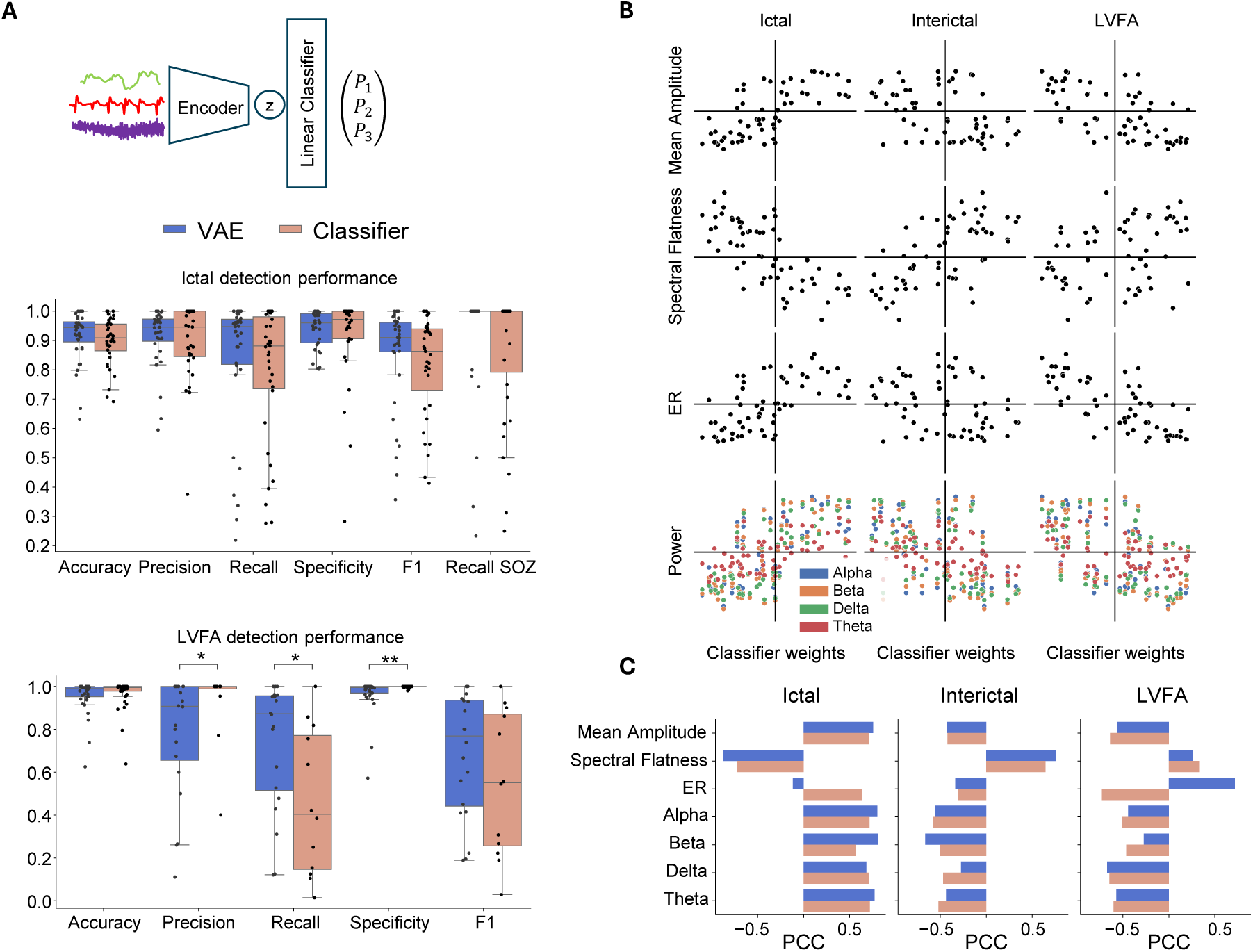
VAE Ablation Study. **A**: Detection performance for ictal and low voltage fast activity (LVFA) comparing full variational autoencoder (VAE) versus encoder-only discriminative model with identical encoder architecture. Points show per-subject performance. Mann-Whitney U tests: *: *p <* 0.05, **: *p <* 0.01. **B**: Linear classifier weight of the encoder only model versus feature correlation scatter plots for each latent dimension, by class. **C**: Pearson correlation coefficients between classifier weights and feature correlations of the VAE and the encoder-only model.

Although Mann-Whitney U tests revealed no statistically significant differences between models for ictal detection, the VAE consistently yielded higher performance across all metrics. Accuracy decreased from 0.92 (VAE) to 0.90 (encoder-only), precision from 0.91 to 0.89, recall from 0.84 to 0.79, specificity from 0.88 to 0.86, and F1 score from 0.85 to 0.81. Notably, recall on seizure onset zone (SOZ) channels decreased from 0.93 to 0.87.

In addition, the VAE achieved significantly more accurate temporal onset marking than the encoder-only model (Mann-Whitney U test, p ¡ 0.0001). For ictal onset detection, the VAE showed a mean time difference of 9.8 seconds (median 5.01 seconds) compared to 13.55 seconds (median 5.9 seconds) for the encoder-only model.

Figure 6A shows contrasting performance patterns for LVFA detection between models. The encoder-only model achieved statistically significant higher precision and specificity compared to the VAE. However, the VAE demonstrated significantly higher recall (p ¡ 0.05, Mann-Whitney U test), while F1 scores are higher for the VAE but with no significant difference. The encoder-only network adopts a conservative detection strategy, resulting in a high false negative rate with an average recall of only 25%. This yields high precision and specificity at the expense of substantially reduced sensitivity. In contrast, the VAE shows more balanced metrics.

To assess whether the reconstruction objective influences latent space organization, we performed the same explainability analysis as in the previous section to the encoder only model. Figure 6B presents scatter plots comparing classifier weights against Spearman correlations between latent dimensions and SEEG features for the classifier. The encoder-only model exhibits similar general patterns to the VAE (Figure 5B). Overall, dimensions correlated with class-relevant features tend to receive higher classifier weights, but with noticeably reduced organization and less clear linear relationships.

To quantify this difference, we computed Pearson correlation coefficients (PCC) between classifier weights and feature correlations for each class-feature combination (Figure 6C). For the ictal class, the VAE shows larger PCC values for mean amplitude, spectral flatness, and power in alpha, beta, and theta frequency bands, indicating stronger alignment between these clinically relevant ictal markers and classifier decision-making. Notably, while the VAe shows negligible correlation with the ER, the encoder-only model exhibits a non-negligible positive correlation. Thus, the VAE classifier is more consistent with the the low-frequency rhythmic characteristics of ictal activity.

The VAE demonstrated higher correlation with spectral flatness for the interictal class, appropriately capturing the broadband, non-rhythmic nature of interictal activity.

The LVFA weights show that while the VAE displays a positive correlation with ER, the encoder-only model shows a negative correlation. Thus, the VAE better aligns with the LVFA’s high-frequency, low-amplitude characteristics. In contrast, the encoder-only model shows slightly larger negative correlation with mean amplitude.

The ablation study reveals that the VAE’s reconstruction objective provides benefits in both performance and explainability. While the encoder-only model achieves comparable classification accuracy, the VAE demonstrates: (1) improved temporal accuracy for ictal onset marking, (2) higher sensitivity for LVFA detection despite lower specificity, and (3) stronger alignment between physiologically interpretable features and classifier weights across all three classes.

These results suggest that the reconstruction task regularizes the latent space organization, encouraging dimensions to encode features in a manner more consistent with clinical understanding of seizure electrophysiology. However, the encoder-only model’s ability to produce similar overall patterns indicates that the linear classifier architecture itself implicitly enforces feature-weight relationships even without explicit reconstruction guidance. The VAE’s reconstruction objective enhances this organization while slightly improving the performance. This suggests that the reconstruction performance may help generalize in challenging detection tasks like LVFA identification and precise temporal onset marking.

## 4 Discussion

Our VAE-based system automatically annotates the onset of both LVFA and ictal activity on individual SEEG channels. In current clinical practice, epileptologists manually review SEEG recordings to delineate SOZs. This process typically involves the identification of seizure events, onset phases and their times through detailed inspection of time and frequency-domain signal features. This is a tedious process, prone to inter-observer variability. A system that can automate this process by detecting ictal activity and LVFA as SOZ indicators, as well as accurately annotating their onset in time, offers a potential solution to streamline this workflow. In a clinical setting, this system could function as a decision support tool by generating preliminary annotations for each channel (adding LVFA and ictal markers). These pre-annotations could then be reviewed and, if necessary, refined by the clinician, thereby preserving clinician oversight while reducing the burden of manual SEEG review.

Unlike black-box neural networks, our Variational Autoencoder learns a latent space that directly correlates latent dimensions with physiologically meaningful SEEG features. After checking the linear classifier’s weights, we can obtain the relevance of each dimension, and therefore feature, per class. Our results have shown that our VAE gives higher importance to features characteristic to each class, such as amplitude, band-power, spectral flatness, and energy ratio. Clinicians can inspect which aspects of the signal drive its predictions. For example, high-amplitude/low-frequency dimensions cause the model to predict an ictal label, whereas dimensions correlated with high-frequency energy ratio and anti-correlated to mean amplitude tip it toward LVFA. Our system presents lower correlations to high frequency features compared to low frequency ones, potentially reflecting the presence of the previously mentioned spectral bias that affects all neural networks. This transparency is crucial when deploying AI in clinical applications, where understanding the system’s outputs can build clinician trust and facilitate its acceptance.

It is worth noting that across the 5-fold validation strategy, optimal postprocessing thresholds lay within a narrow range, with the lowest being 6.18 and the highest 6.54, demonstrating the robustness of the postprocessing algorithm across multiple subjects. Although the segment level classification, especially between the ictal and LVFA classes, indicate a high degree of misclassifications, our post-processing algorithm mitigates their impact at the channel level. We obtained high ictal detection metrics for the majority of subjects, with enhanced performance on channels labeled as SOZ, indicating that most of misclassifications happen for recordings outside of the SOZ that could be deemed ambiguous.

Although the system demonstrates strong detection performance for the majority of subjects, there is a subset that exhibits markedly reduced accuracy. Analysis of these cases revealed common trends that cause the system to fail. As shown in Supplementary material figure S2, very short ictal periods (lower than 10 seconds) are less likely to be detected because almost all segments need to be assigned a high ictal probability for the AUC to surpass the threshold. In relation to this, we have identified common causes contributing to low ictal prediction probabilities. ictal segments presenting lower amplitude and/or higher frequency than the average caused the output ictal probability to decrease. In some cases, this shift favors the LVFA probability. This behavior is exemplified in figure S3. Moreover, a significant amount of the failures in the ictal detection can be attributed to the spatial resolution of SEEG recordings. Due to the inter-electrode spacing of 1–1.5 mm typically found in SEEG leads, adjacent bipolar channels in the same SEEG lead may present subtle electrographic variations. Therefore, the transition between bipolar channels within and outside the SOZ can introduce ambiguity alongside the SEEG lead. This increases the likelihood of misclassification between interictal and ictal activity and decreases the temporal accuracy of ictal onset marker in our system, as demonstrated in Supplementary Figure S4. The same scenario is also found, to a lesser extent, in the LVFA class. Minor variations in the spectral content across neighboring bipolar channels in the same lead can be seen, causing misclassifications between the LVFA and interictal phases.

LVFA detection achieved 74% recall, lower than ictal detection (86%) warranting detailed analysis of mislabeling sources. We identify four primary challenges: (1) Spectral bias in neural networks: High-frequency LVFA components are systematically under-reconstructed by the VAE (Figure 2A), a fundamental limitation of gradient-based optimization that preferentially learns low-frequency patterns. This makes LVFA harder to distinguish from baseline activity. (2) Severe class imbalance: Despite data augmentation, LVFA segments represent only 3.7% of the dataset (Table 2), limiting the model’s exposure to these patterns. (3) Artifact sensitivity: a large number of LVFA misclassifications are caused by the presence of high frequency noise in the interictal phase, as in the example shown in the Supplementary Figure S5, indicating the lower robustness of the system when annotating LVFA compared to the ictal phase. (4) Ictal Annotation dependency: By clinical definition, LVFA markers require subsequent ictal activity on the same channel. By design we condition LVFA detection on ictal presence, failed ictal detection therefore causes secondary LVFA annotation failures (see example presented in Supplementary Figure S6), even when LVFA patterns are correctly identified at the segment level.

To the best of our knowledge, automated detection of LVFA has been addressed in only a handful of prior studies, none of which has a task formulation directly comparable to ours. Gnatkovsky et al.^44^ proposed a per-channel detector for fast activity at seizure onset based on band-passed energy and a signal-flattening index, which produces explicit onset markers but does not report quantitative detection or temporal-accuracy metrics against expert LVFA annotations. Makaram et al.^45^ and Mohagheghian et al.^46^ trained an SVM and a Random Forest, respectively, on hand-crafted features to classify LVFA among other seizure onset patterns, reporting LVFA F-scores of 90.8 % and 81.4 %. Both, however, perform classification on expert-segmented onset segments restricted to SOZ channels rather than detection of LVFA as a temporal event on all channels of a continuous SEEG recording. These fundamental differences in task formulation, data, and evaluation protocol do not allow for a direct quantitative comparison with our system.

Most scientific work in SEEG analysis has focused around binary seizure detection.^16, 17^ Desai et al.,^26^ using transfer learning on implanted iEEG, reported seizure onset time prediction with a median absolute error of 3.1 seconds after fine-tuning, compared with 4.4–4.7 seconds in less optimized variants. Wong et al.^47^ reported strong segment-level seizure (ictal activity) detection on scalp EEG with AUC 0.93, sensitivity 0.82, and specificity 0.88, which is comparable to our results for ictal detection. Furthermore, their temporal resolution to detect seizures is of 30 seconds compared to our 0.5 seconds. Kharbouch et al.^48^ detected 97% of seizures across 875 hours of iEEG with a median detection delay of 5 seconds and a median false alarm rate of 0.6/day. This latency is shorter than our median ictal onset lag (9.8 seconds) but shorter than our median LVFA onset lag (2.0 seconds). Ojemann et al.^49^ provides an unsupervised seizure annotation tool by modeling patient specific neural dynamics using LSTMs, detecting seizures as deviations from interictal behavior. The method achieved AUROC values around 0.87–0.89. Wang et al.^50^ evaluated 1D CNNs on the SWEC-ETHZ iEEG dataset, both at a segment-based level, where a 90.09% sensitivity, 99.81% specificity and 99.73% accuracy were obtained. In the event-based level, 97.52% sensitivity, and mean 13.2 second latency were attained.

Overall, while some existing systems achieve similar or marginally better results, our work stands out by combining features that other research have targeted individually into a single tool, such as strong seizure detection performance, clinically relevant ictal and LVFA phase differentiation, and an interpretability component through structured latent space analysis. To the best of our knowledge, this is the only work that incorporates these features on iEEG data.

Although seizure detection was not our primary objective, we evaluated the system’s performance as an event detector and obtained a seizure detection rate of 99.1% with a false positive rate of 1% and a false negative rate of 4.5%. Compared with prior episode-level seizure detectors, our VAE-based SEEG system is clearly competitive while addressing a more clinically specific task. Gu et al.^51^ reported 79.6% sensitivity on CHB-MIT and 78.0%, Jeong et al.^52^ reached AUROC 0.98 on pediatric clinical EEG, and Gabeff et al.^53^ reported F1 = 0.873 with roughly 90% seizure detection on iEEG. In comparison, our method places in the same performance range, while also incorporating the interpretabiltiy component.

### The role of the reconstruction objective (compression bias)

The ablation study demonstrates that the VAE provides modest but consistent performance advantages over the encoder-only model, particularly in temporal accuracy and LVFA detection. While ictal detection metrics showed no significant differences, the VAE consistently outperformed across all measures. For LVFA detection, the VAE achieved more balanced performance with higher recall (74% vs. 25%), whereas the encoder-only model’s focus on pure classification resulted in a conservative strategy that missed the majority of LVFA events. These results align with multi-task learning literature demonstrating that auxiliary reconstruction objectives can improve primary task performance by encouraging more complete feature representations and providing regularization.^54–56^ The VAE’s reconstruction objective may promote better generalization by forcing the encoder to preserve features necessary for signal reconstruction rather than only those minimally sufficient for classification, though a larger dataset would be needed to confirm this hypothesis.

A purely discriminative encoder is optimized only for label prediction and therefore need retain only those features that separate the training classes. When spurious or shortcut features correlate with the labels in the training distribution, such a model may exploit them rather than learning features that transfer across subjects, sessions, or acquisition conditions.^57^ Adding a reconstruction objective changes this inductive bias. The encoder is no longer trained solely to support the classification boundary, but also to preserve information about the input signal that enables its reconstruction. In the variational case, this yields a latent-variable generative model in which the encoder approximates *q_ϕ_*(*z|x*) and the decoder defines *p_θ_*(*x|z*).^58, 59^ This does not guarantee recovery of the true physiological generative factors: the latent space can still encode nuisance structure or artefacts if these help reconstruction.^60^ However, under a constrained latent channel, limited decoder capacity, and appropriate regularisation, the auxiliary reconstruction task biases the representation toward compact factors that explain systematic structure in the signal rather than only label-correlated discriminative cues. This is the standard multi-task-learning rationale: an auxiliary task can act as an inductive bias that improves generalisation when it shares structure with the target task.^61, 62^

The information-theoretic interpretation is a rate–distortion one. In a VAE, the expected KL term controls the coding rate of the latent channel and is closely related to an upper bound on *I_q_*(*X*; *Z*), while the reconstruction term penalises distortion between *x* and its reconstruction. The ELBO thus trades off compression of the latent code against preservation of information needed to explain the input, connecting VAEs to information-bottleneck-style representation learning^63^ and to the broader principle that more compactly describable models tend to generalize better.^64^

In our setting, the SEEG signal is forced through a narrow stochastic bottleneck while still being reconstructable, biasing the classifier toward compact latent structure in the signal. Empirically, this is consistent with the observed alignment of latent dimensions with physiologically interpretable spectral and amplitude features, and with the improved behaviour on the ambiguous LVFA class, where the label-only model collapses to a conservative high-precision, low-recall strategy that misses most LVFA events.

Consistent with this interpretation, the VAE showed substantially stronger alignment between latent dimensions and physiologically relevant features, suggesting that reconstruction regularizes the latent space toward more coherent feature encoding. Interestingly, the encoder-only model exhibited similar organizational patterns, likely due to the linear classifier architecture imposing inherent constraints, though these patterns were considerably weaker and less physiologically consistent than in the VAE.

## Limitations

While our VAE-based pipeline showcases strong performance in seizure phases annotation and detection, we have identified several main lines of limitations and potential improvements. First, despite ictal onset and presence information being provided by clinicians in both datasets, no information on the LVFA was available. To include LVFA onset phase, we adopted a systematic, but ultimately subjective protocol to label LVFA on individual channels. Despite this fact and the limited number of examples, our results are promising and show the potential of the proposed approach to accurately detect LVFA. Furthermore, future studies could include more clinically relevant seizure onset patterns besides the common LVFA, such as rhythmic spiking or warp theta-band bursts.^65^ To address all this in future studies, a larger dataset as well as expert annotations of all these seizure onset patterns would be needed.

Second, the system only outputs categorical markers per channel. However, the predicted probabilities per segment could be used to provide confidence scores to clinicians, allowing them to focus their review on low-confidence predictions. Explainable AI techniques such as confidence scores have shown to improve trust.^66, 67^ Furthermore, our current approach only considers the information of each channel for its annotation. Epileptic activity, however, is known to propagate across brain networks. By incorporating multichannel context, for instance using cross-channel attention,^47, 68^ we could hypothetically capture propagation patterns from SOZ zones, potentially improving detection performance and providing physiological information to the output confidence scores in propagation zones.

Additionally, further work should be aimed at mitigating the apparent spectral bias in the current system. Such behavior could be mitigated by incorporating loss terms that heavily target high frequency information, such as the spectral mean squared error or, with the addition of a small neural network acting as a discriminator, therefore introducing the GAN training objective, the adversarial loss.^69^

Lastly, an important limitation of this study is the relatively small dataset available for training and evaluating the system. A larger dataset would greatly benefit further work. In addition, a systematic architecture search or systematic hyperparameter optimization on larger datasets could identify more efficient or effective architectures. Our architecture follows established CNN-LSTM design patterns for physiological signals, but alternative configurations may achieve equal or superior performance.

## 5 Conclusions

We have demonstrated that a Variational Autoencoder–based architecture can deliver both accurate and explainable annotations of SEEG data. By encoding 2-second segments into a physiologically meaningful latent space and coupling it with a simple linear classifier, our system achieves channel-level ictal recall above 84% (rising to over 91% in SOZ channels) and LVFA recall of 74%, while maintaining median temporal error under 10 seconds for ictal onsets and under 1 second for LVFA. As a seizure detector, our system provides a 99% detection recall with a 4.5% false negative rate.

Beyond its performance, our framework is designed to accelerate clinician SEEG review times. By pre-annotating channels with markers for ictal onset and LVFA, the system can dramatically reduce the hours of manual labor required, prioritize the review of likely seizure onset zones, and potentially minimize inter-observer variability, acting as a decision support system.

Finally, the correlation between the latent space and physiological SEEG features, coupled with the importance the classifier gives to each class, makes the system a potential candidate for real-world clinical translation. The possibility of inspecting which latent dimensions play a role in classification, for instance, low-frequency rhythmicity vs. high-frequency bursts, driving decisions, will ease its acceptance in clinical settings. In summary, our work establishes a robust, explainable, and clinically based platform for automated SEEG annotation that can support clinicians in epilepsy surgery planning.

## Acknowledgements

This work has received funding from the European Research Council (ERC) under the European Union’s Horizon 2020 research and innovation programme (Grant Agreement No. 855109; ERC-SyG 2019) and from FET under the European Union’s Horizon 2020 research and innovation programme (Grant Agreement No. 101017716).

## Data Availability

The data supporting the findings of this study are derived from the following sources:

- The HUP iEEG dataset is publicly available through OpenNeuro (https://openneuro.org/datasets/ds004100/versions/1.1.1).
- The GALVANI project data were provided by the Clinical Physiology Department, INSERM, UMR 1106, and Timone University Hospital under license for the current study. This dataset is not publicly available as it’s a result of two clinical trials (NCT04782869, NCT05250713)

## Ethical Standards

This work has used human data already generated, either publically available or as a result from clinical trials. This study did not generate any new human data and all data was de-identified by the provider sources.

