## Supplementary Materials for "Variational autoencoder for Explainable seizure onset phases detection"

### Supplementary materials for Variational autoencoder for interpretable seizure onset phases detection

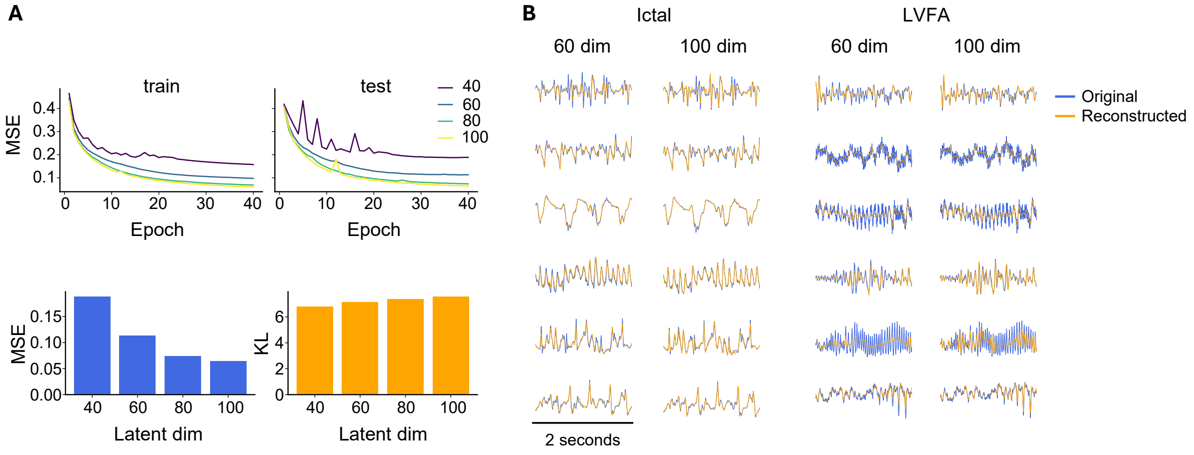


Figure S1: A) Systematic evaluation of latent dimension size impact on reconstruction quality. Upper panels: Training and test reconstruction loss (variance-normalized mean squared error, MSE) across epochs for latent dimensions 40, 60, 80, and 100. Bottom panels: Final test set MSE and KL divergence after training completion. Reconstruction loss decreases monotonically with increasing latent dimensions but with diminishing returns beyond 60 dimensions. 60, 80, and 100 dimensions achieve comparable performance with minimal overfitting (training and test curves remain closely aligned). KL divergence increases slightly with dimension size, reflecting the expanding latent space, but remains well-controlled across all configurations. These models were trained using only reconstruction and KL divergence losses, without the classification objective, to isolate the effect of latent dimension size on reconstruction capacity. B) Representative reconstruction examples comparing 60 and 100 dimensions for ictal and LVFA segments. For the ictal fragments both latent space sizes preserve the dominant amplitude modulation and temporal dynamics. In contrast, high-frequency, low-amplitude LVFA patterns remain challenging to reconstruct at both 60 and 100 dimensions. While 100 dimensions provides marginally better high-frequency preservation, the reconstruction quality improvement is modest compared to the substantial increase in model complexity.

*

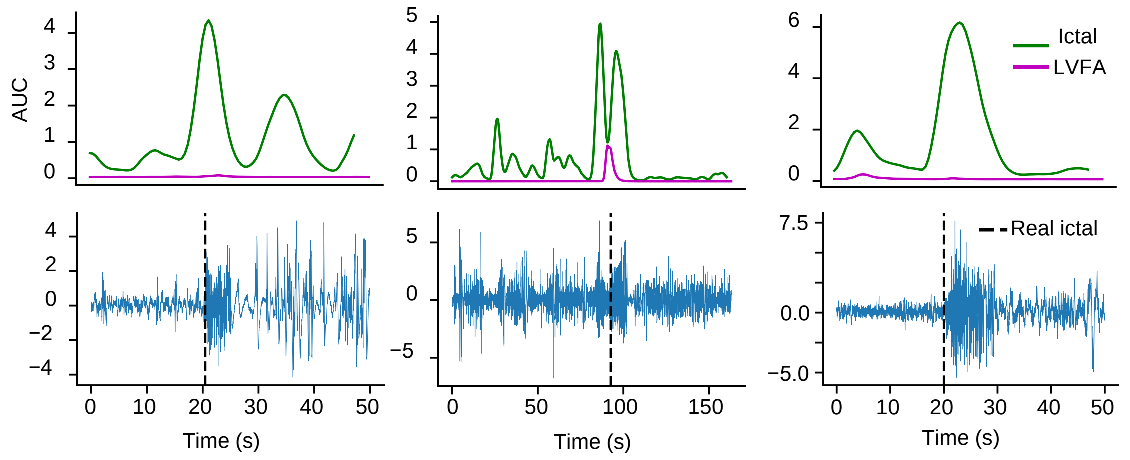


Figure S2: Examples of bipolar SEEG channels from different subjects. From top to bottom, the plots indicate the AUC signal produced by the ictal/LVFA probabilities of the fragments composing the channel and the SEEG channel with the real and predicted markers, if any. A very short ictal period can cause the AUC to not reach detection threshold unless all fragments have a high ictal probability, leading to ictal detection failure.

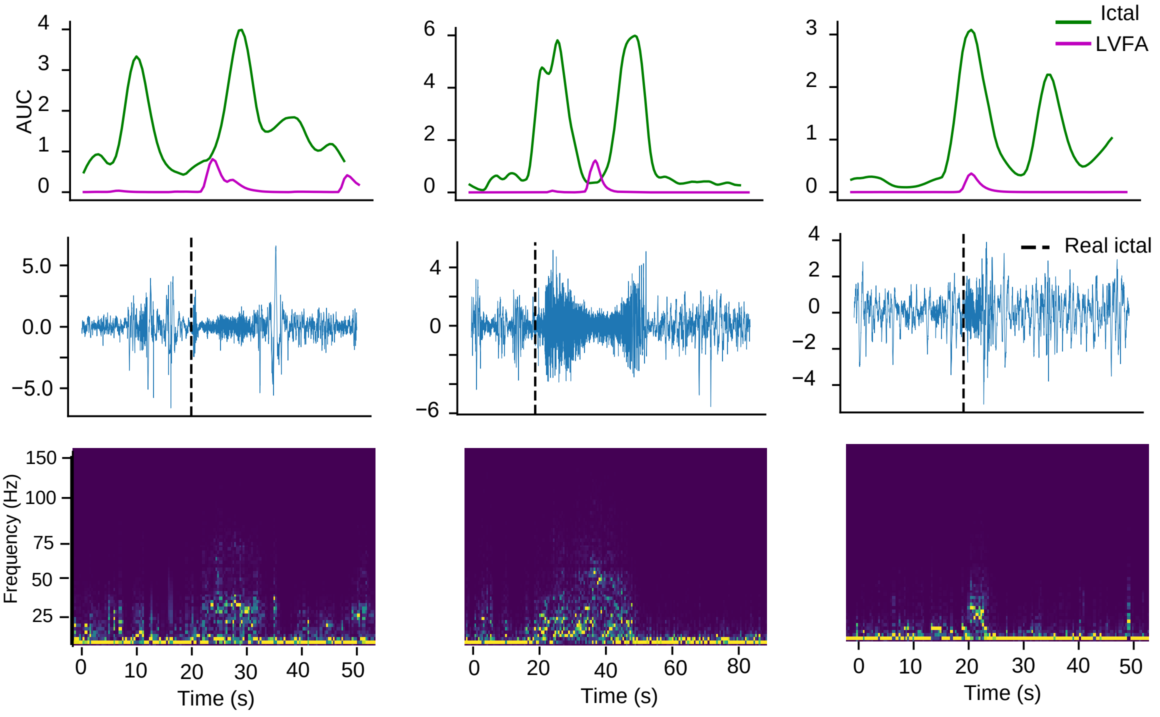


Figure S3: Examples of bipolar SEEG channels from different subjects. From top to bottom, the plots indicate the AUC signal produced by the ictal/LVFA probabilities of the fragments composing the channel, the SEEG channel with the real and predicted markers, if any, and the spectrogram of the SEEG channel. Channels with an ictal activity at low amplitude and higher frequency than the average ictal activity cause the ictal AUC to be lower, therefore not reaching the detection threshold. Additionally, the LVFA AUC increases on the same period due to confusion between the two classes.

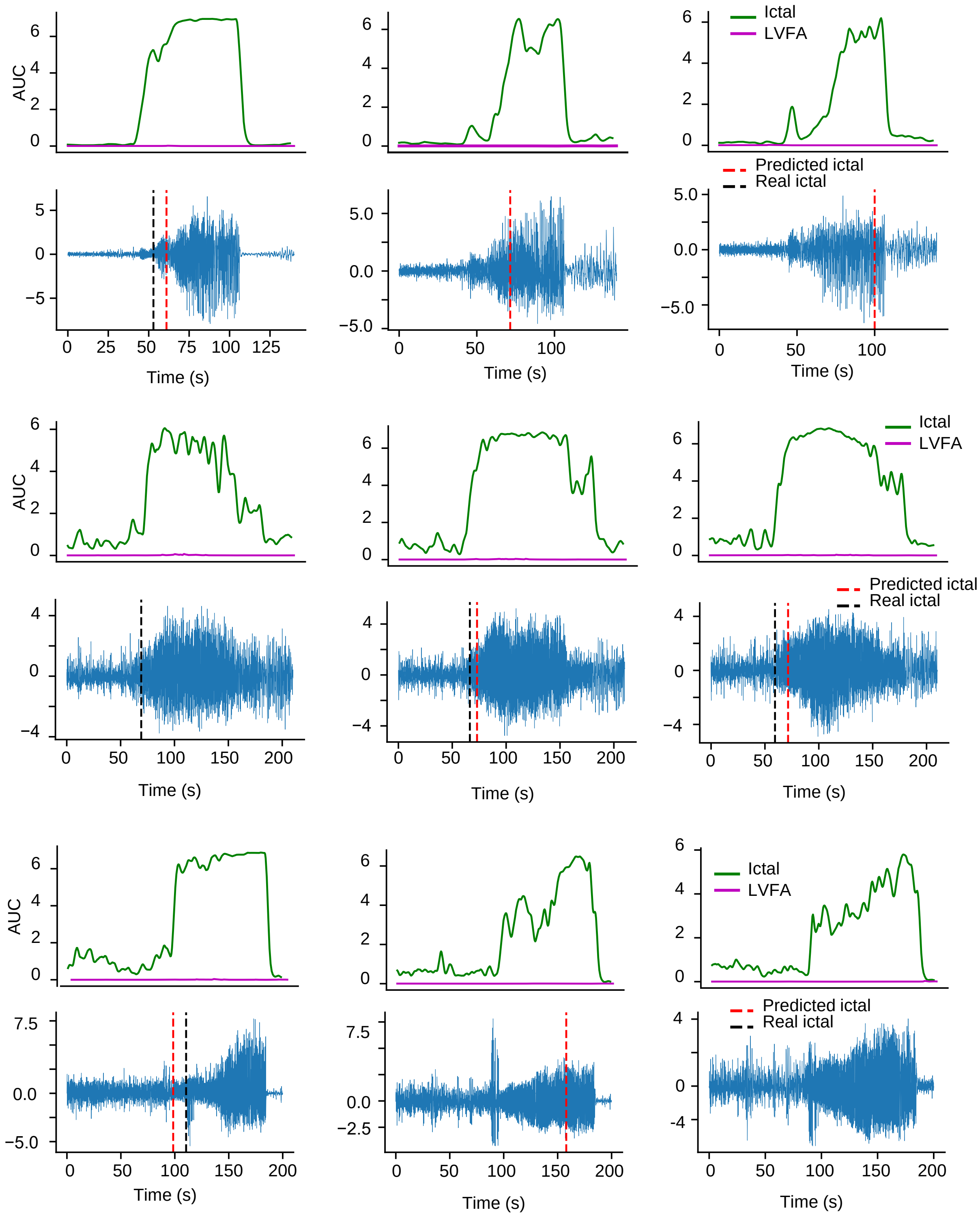


Figure S4: Examples of contiguous bipolar SEEG channels along the same SEEG leads from different subjects. From top to bottom, the plots indicate the AUC signal produced by the ictal/LVFA probabilities of the fragments composing the channel and the SEEG channel with the real and predicted markers, if any. Due to an inter-electrode spacing of 1-1.5mm, there are brief differences between adjacent electrodes, causing ambiguity that causes difficulty to the system when differentiating between interictal and ictal activity, particularly as the SEEG lead progresses out of the SOZ.

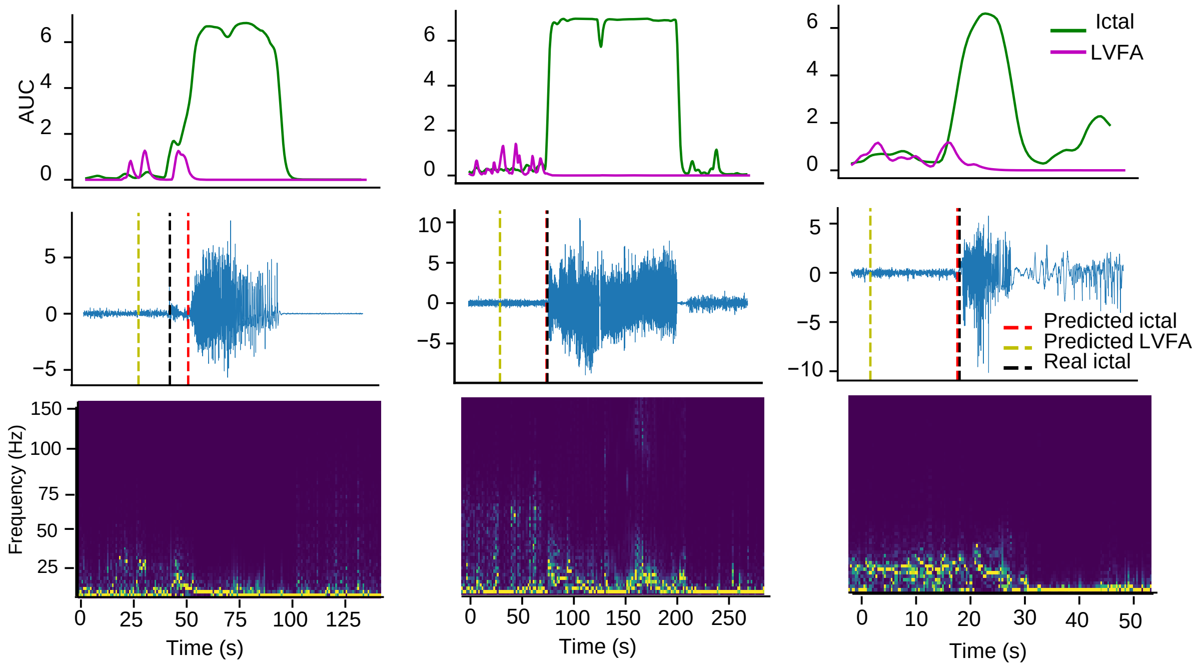


Figure S5: Examples of contiguous bipolar SEEG channels from different subjects.From top to bottom, the plots indicate the AUC signal produced by the ictal/LVFA probabilities of the fragments composing the channel, the SEEG channel with the real and predicted markers, if any, and the spectrogram of the SEEG channel. The presence of unrelated high frequency activity in the signal causes our system to produce false positive predictions for the LVFA onset annotations.

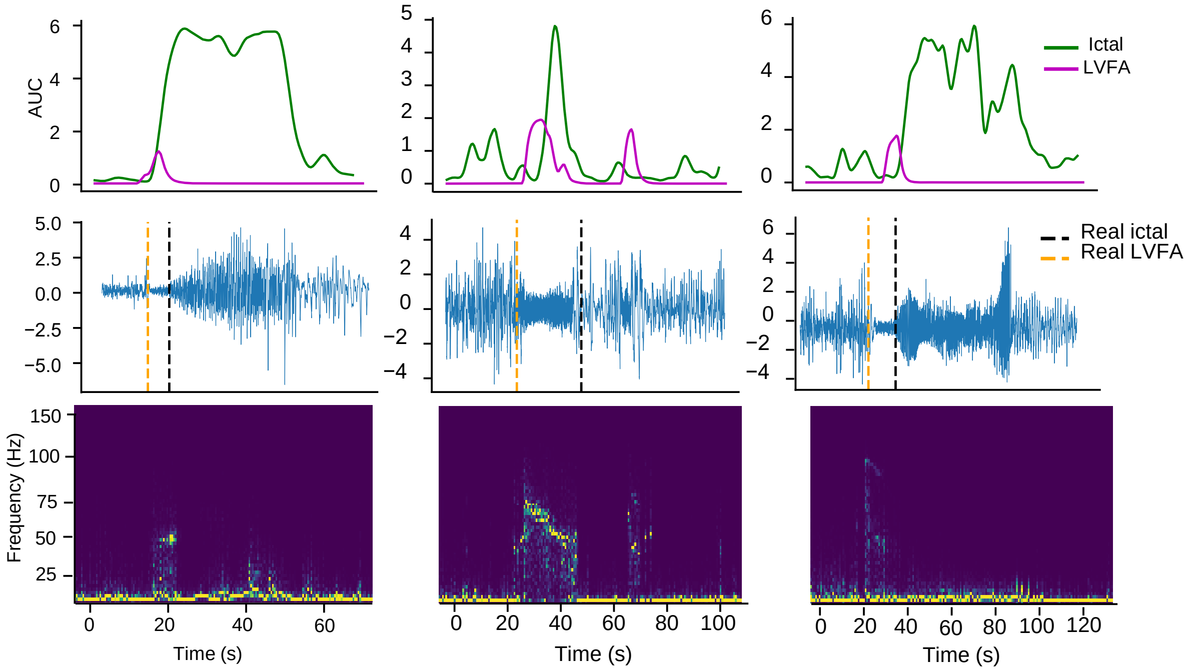


Figure S6: Examples of contiguous bipolar SEEG channels from different subjects. From top to bottom, the plots indicate the AUC signal produced by the ictal/LVFA probabilities of the fragments composing the channel, the SEEG channel with the real and predicted markers, if any, and the spectrogram of the SEEG channel. The AUC plots show that despite a good identification of the LVFA phase. The ictal AUC does not reach the threshold and therefore ictal detection fails. As a result, given our design choice to only consider predicted LVFA onset markers if there is also a predicted ictal onset marker, the LVFA detection fails.

| Subject | age | gender | age at onset | Epilepsy type | MRI | Seizures | Channels | SOZ channels | LVFA channels |
|---|---|---|---|---|---|---|---|---|---|
| **GAL01** | 21 | F | 14 | Parietal | Lesional | 3 | 142 | 8 | 4 |
| **GAL02** | 23 | F | 9 | Occipital | Non-Lesional | 3 | 171 | 6 | 3 |
| **GAL03** | 24 | F | 10 | Temporal | Lesional | 4 | 83 | 16 | 35 |
| **GAL04** | 30 | M | 11 | Temporal | Non-Lesional | 2 | 191 | 15 | 0 |
| **GAL05** | 49 | M | n/a | Temporal/MTL | Non-Lesional | 1 | 166 | 6 | 8 |
| **GAL06** | 32 | F | | Motor | Lesional | 2 | 218 | 17 | 0 |
| **GAL07** | 35 | M | 19 | Temporal | Non-Lesional | 1 | 99 | 12 | 0 |
| **GAL08** | 15 | M | n/a | Motor | Non-Lesional | 1 | 165 | 22 | 2 |
| **GAL09** | 49 | M | n/a | Temporal | Non-Lesional | 2 | 161 | 3 | 0 |
| **GAL10** | 27 | M | 10 | Temporal | Non-Lesional | 3 | 166 | 11 | 75 |
| **GAL11** | 33 | M | 12 | Frontal | Non-Lesional | 3 | 135 | 11 | 10 |
| **GAL12** | 44 | M | 12 | Frontal | Non-Lesional | 3 | 153 | 6 | 37 |
| **GAL13** | 44 | F | 34 | Temporal/Insular | Non-Lesional | 1 | 140 | 13 | 41 |
| **HUP060** | 42 | F | 12 | Frontal | n/a | 3 | 36 | 3 | 0 |
| **HUP116** | 59 | F | 42 | MTL | Lesional | 2 | 37 | 6 | 0 |
| **HUP117** | 39 | M | 12 | Temporal | Lesional | 1 | 35 | 6 | 0 |
| **HUP133** | 52 | F | 47 | MTL | Non-Lesional | 5 | 64 | 4 | 6 |
| **HUP134** | 32 | M | 8 | Frontal | Lesional | 1 | 71 | 5 | 0 |
| **HUP135** | 37 | M | 34 | MTL | Non-Lesional | 2 | 92 | 2 | 0 |
| **HUP138** | 38 | M | 29 | MTL | Lesional | 4 | 90 | 3 | 0 |
| **HUP140** | 47 | F | 26 | MTL | Non-Lesional | 3 | 73 | 4 | 0 |
| **HUP142** | 30 | M | 15 | MTL | Lesional | 3 | 99 | 15 | 0 |
| **HUP144** | 31 | M | 5 | Temporal | Lesional | 5 | 96 | 5 | 15 |
| **HUP146** | 16 | M | 4 | Temporal | Non-Lesional | 3 | 86 | 11 | 4 |
| **HUP148** | 23 | M | 16 | Temporal | Lesional | 4 | 90 | 25 | 9 |
| **HUP150** | 17 | M | 4 | Insular | Lesional | 5 | 79 | 8 | 0 |
| **HUP151** | 33 | M | 6 | MFL | Non-Lesional | 5 | 154 | 2 | 4 |
| **HUP157** | 25 | M | 16 | MTL | Non-Lesional | 5 | 145 | 13 | 33 |
| **HUP158** | 32 | M | 7 | Insular | Non-Lesional | 1 | 158 | 4 | 0 |
| **HUP160** | 45 | F | 15 | Temporal | Non-Lesional | 3 | 92 | 21 | 0 |
| **HUP163** | 42 | F | 32 | MTL | Non-Lesional | 3 | 143 | 6 | 1 |
| **HUP164** | 34 | F | 14 | MTL | Lesional | 3 | 160 | 9 | 10 |
| **HUP166** | 26 | M | 4 | Temporal | Lesional | 2 | 145 | 5 | 0 |
| **HUP180** | 28 | F | 2 | Frontal | Lesional | 5 | 101 | 8 | 0 |
| **HUP187** | 25 | M | 18 | MTL | Non-Lesional | 5 | 78 | 14 | 0 |
| **HUP188** | 24 | F | 9 | Frontal | Lesional | 5 | 146 | 18 | 0 |
| **HUP190** | 25 | M | 12 | MTL | Non-Lesional | 3 | 127 | 12 | 13 |

Table S1: Subject-level demographic and clinical characteristics. Age in years at time of SEEG monitoring. Epilepsy types: MTL = mesial temporal lobe epilepsy. Channels: bipolar channel count after preprocessing. SOZ Size = number of channels clinically identified as seizure onset zone. LVFA channels indicates the number of channels in which low-voltage fast activity was annotated in any seizure for that subject. Dataset Source: subjects whose ID contains ”HUP” are Hospital of University of Pennsylvania public dataset, and subjects whose ID contains ”GAL” are from GALVANI project.
